# EGRET: Edge Aggregated Graph Attention Networks and Transfer Learning Improve Protein-Protein Interaction Site Prediction

**DOI:** 10.1101/2020.11.07.372466

**Authors:** Sazan Mahbub, Md Shamsuzzoha Bayzid

**Affiliations:** Department of Computer Science and Engineering, Bangladesh University of Engineering and Technology, Dhaka-1205, Bangladesh; Department of Computer Science, University of Maryland, College Park, Maryland 20742, USA

**Keywords:** Protein-Protein Interaction Sites, Deep Learning, Graph Neural Network, Edge Aggregation

## Abstract

**Motivation:** Protein-protein interactions are central to most biological processes. However, reliable identification of protein-protein interaction (PPI) sites using conventional experimental methods is slow and expensive. Therefore, great efforts are being put into computational methods to identify PPI sites.

**Results:** We present EGRET, a highly accurate deep learning based method for PPI site prediction, where we have used an edge aggregated graph attention network to effectively leverage the structural information. We, for the first time, have used transfer learning in PPI site prediction. Our proposed edge aggregated network, together with transfer learning, has achieved notable improvement over the best alternate methods. Furthermore, we systematically investigated EGRET’s network behavior to provide insights about the causes of its decisions.

**Availability:** EGRET is freely available as an open source project at https://github.com/Sazan-Mahbub/EGRET.

**Key Points:**

- We present a comprehensive assessment of a compendium of computational protocols to solve an important problem in computational proteomics.
- We present a highly accurate deep learning method, EGRET, for Protein-Protein Interaction (PPI) site prediction for isolated proteins.
- We have used an edge aggregated graph attention network to effectively capture the structural information for PPI site prediction.
- We, for the first time, present a successful utilization of transfer-learning from pretrained transformer-like models in PPI site prediction.

## 1 Introduction

Proteins are responsible of various functions in cells, but carrying out many of these function requires the interaction of more than one protein molecules [1]. This makes protein-protein interaction (PPI) one of the key elements for understanding underlying biological processes, including functions of the cells [2, 3], PPI networks [1, 4], disease mechanisms [5, 6], as well as for designing and developing novel therapeutics [7, 8].

Protein-protein interaction (PPI) sites are the interfacial residues of a protein that interact with other protein molecules. Several wet-lab methods, including two-hybrid screening and affinity purification coupled to mass spectrometry are usually used to identify PPI sites [9, 10, 11]. However, the experimental determination of PPI sites is costly and time- and labour-intensive [12, 13, 14]. Thus, highly accurate computational prediction methods can be a useful guide for and complement to genetic and biochemical experiments. Therefore, in the last two decades, computational approaches have emerged as an important means of predicting PPI sites [13, 15, 16, 17]. These computational methods can be roughly divided into three categories [18]: (1) Protein-protein docking and modeling [19, 20], (2) Structure-based methods [15, 16, 21, 22, 23, 24], and (3) Sequence based methods [18, 25, 26, 27, 28, 29]. Docking and structure based methods, unlike the sequence-based methods, leverage the structural information of the protein molecules.

There are two major areas in protein-protein interaction sites prediction (PPISP). One is pair-wise interaction sites prediction for predicting interfacial residues of a pair of proteins, which is related to the docking of two proteins [30, 31, 32]. The second prediction problem – and the one addressed in this study – is the prediction of putative interaction sites upon the surface of an isolated protein, known to be involved in protein-protein interactions, but where the structure of the partner or complex is not known [33]. The absence of any information about the partner proteins makes the latter problem relatively more difficult and challenging [31, 34].

In order to predict the interfacial residues of a single (isolated) protein, most of the recent computational methods have applied various machine learning (ML) algorithms [15, 16, 18, 21, 25, 28, 29, 35, 36]. Many studies have shown the importance of using local contextual features for predicting interfacing residues [15, 18, 37, 38, 39], which is usually encoded by a sliding window with a fixed size. However, similar to other residue-level prediction problems, such as secondary structure, backbone torsion angle, relative accessible surface area [40, 41], information about the *long-range interactions* between the residues that are not sequentially closer but within a close proximity in three-dimensional Euclidean space is also very crucial for PPI sites prediction. DeepPPISP [15] addressed this issue with global features, extracted using two-dimensional convolutional neural networks. But such global feature extraction method generates the same global feature representation for all the residues of a protein, and thus lacks the ability to learn various suitable functions for different residues, and subsequently may not effectively encode long-range interactions between different residues.

Position-Specific Scoring Matrix (PSSM) is one of the most useful features for the prediction of PPI sites [15], which is also widely used by many state-of-the-art methods [15, 16, 25]. Other features, such as primary sequence and secondary structure information, alone do not yield good predictions when PSSM features are not used [15]. Generation of PSSM using PSI-BLAST [42], however, is a time-consuming process, which [26] pointed out as a major bottleneck of these methods. Hence, identifying the feature-sets, which are less computationally demanding yet effective for highly accurate prediction of PPI sites is of great interest. The recent advancement in Natural Language Processing (NLP) can contribute in this direction as a trained language model can extract features to use as input for a subsequently trained supervised model through transfer-learning [43]. In a very recent study, [43] developed ProtTrans, which provides an outstanding model for protein pretraining. They showed that a new set of features, generated by pre-trained transformer-like models are capable of performing very well while taking significantly less time compared to PSSM. They trained two auto regression language models (Transformer-XL [44] and XLNet [45]) and two auto encoding models (BERT [46] and Albert [47]) on data containing up to 393 billion amino acids from 2.1 billion protein sequences in a self-supervised manner, considering each residue as a “word” (similar to language modeling in natural language processing [46]). In a similar study, [48] showed that “attention scores” in some of the attention matrices of such pretrained transformer-like models correlate in various degrees with protein-attributes, including contact maps and certain types of binding sites.

Most of the existing methods for predicting PPI use features derived from the primary sequences of the proteins [15, 16, 18, 21, 25, 26, 28, 29, 36, 49]. However, using only primary sequence based features may limit the capabilities of the methods to achieve higher accuracy [21]. Thus, several methods have leveraged features derived from structural data (generally from the PDB files [50]), including three state and eight state secondary structures [15, 16], the level of surface exposure to solvent [21, 22], local surface region in a query protein structure [23], etc. Effective utilization of the three-dimensional structural information has the potential to increase the performance of PPI sites prediction methods. Graph neural networks (GNN) have been emerged as an effective tool for encoding structural information [30, 51, 52, 53, 54]. However, although GNN based architecture has been applied to pairwise binding site prediction [30, 54], it has not been used for predicting the binding sites of a single protein. Moreover, unlike methods that may not appropriately encode residue-specific information about long-range interactions as they learn a single global feature representation for all the residues (e.g., global representation of proteins by [15]), GNNs have the potential to effectively encode global features involving any specific residue, by learning a suitable function for *a particular residue and its close proximity neighbours*.

Among various GNN based architectures, Graph Attention Network (GAT) was proved to be very effective in protein-interaction network related problem [52]. GAT uses attention mechanism [55, 56] in the node-level aggregation process, which helps it perform a weighted aggregation. But the originally proposed architecture of GAT does not consider the features of the edges, either during *the aggregation process* or during *the calculation of the attention score*. Therefore, GAT lacks the ability to utilize the rich structural information that might have been encoded in the edge-features.

In this paper we leverage a variant of GAT, using both node and edge features during the computation of attention scores [57] which we call **E**dge Aggregated **GR**aph Attention N**ET**work (EGRET), for predicting interaction sites of a single (isolated) protein with known structure. Unlike GAT, EGRET is expected to effectively leverage the structural information encoded in the edge-features during both the aggregation and attention score calculation phases. We also present a successful utilization of transfer-learning from pretrained transformer-like models by [43] in PPI sites prediction. Combined with the transfer-learning, our proposed EGRET architecture improves upon the best alternate methods on the widely used benchmark dataset assembled by [15].

## 2 Approach

### 2.1 Feature Representation

#### 2.1.1 Graph representation of proteins

The proposed model EGRET is a graph neural network based architecture, and we represent the three-dimensional structure of each protein *P* in our dataset as a directed *k*-nearest neighbor graph *G* [58]. The set *V* (*G*) of the nodes in the graph *G* is the set of the amino-acid residues of a protein *P*. Let 𝒩_*i*_ be *the neighborhood* of the node (residue) *i* ∈ *V* (*G*) comprising its *k* nearest neighbors (i.e., |𝒩_*i*_| = *k*), and *i* be the *center of this neighborhood*. Here *i* is connected by directed edges to all the nodes in 𝒩_*i*_. These neighbours of *i* are selected by sorting all other nodes based on their distances from *i*, and then taking the nearest *k* nodes, where *k* is a hyperparameter of our method. Inspired by the success of pair-wise protein-protein interaction prediction by [30], the distance between any two nodes (residues) is calculated by averaging the distances between their atoms (using the atom coordinates from the PDB files [50]). As each residue is represented as a node in our graph representation, we use the terms ‘residue’ and ‘node’ interchangeably for convenience.

#### 2.1.2 Node-level feature representation

Each node *i* ∈ *V* (*G*) (representing the *i*-the residue in a protein sequence) is represented by a feature vector. EGRET takes as input a sequence *X* = {*X*_1_, *X*_2_, *X*_3_, …, *X*_*N*_} of aminoacid residues of the protein *P*, where *X*_*i*_ is the one-letter notation [59] of the *i*-th residue and *N* is the total number of residues in *P*. *X* is passed through the embedding generation pipeline developed by [43]. Although we have used ProtBERT for our experiments since it was shown to achieve superior performance compared to other methods on residue-level classification tasks (e.g. secondary strucuture prediction), EGRET is agnostic to the pretrained language models available in ProtTrans [43], which means that other appropriate language models in ProtTrans can also be used in our method. ProtBERT generates a sequence of embedding vectors 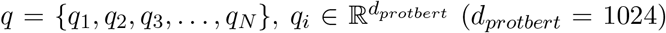, where *q*_*i*_ is used as the node-level feature vector of node *i*.

#### 2.1.3 Edge-level feature representation

In the directed graph representation *G* of a protein *P*, the edge features of an edge *E*_*ji*_ (from node *j* to node *i*) in *G* is denoted by *ξ*_*ji*_, where 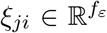 and *f*_*ε*_ is the number of features of the edge. We used the following two features (i.e., *f*_*ε*_ = 2) as edge-features: (1) distance *D*_*ij*_ between the residues *i* and *j*, which is calculated by taking the average of the distances between their atoms, and (2) relative orientation *θ*_*ij*_ of the residues *i* and *j*, which is calculated as the absolute value of the angle between the surface-normals of the planes of these two residues that go through the alpha Carbon atom (*C*_*α*_), Carbon atom of the Carboxyl group, and Nitrogen atom of the Amino group of each of the residues. We standardize both these features across all the training samples.

### 2.2 Architecture of EGRET

The architecture of EGRET can be split into three separate discussions: 1) the architecture of local feature extractor, 2) the architecture of our proposed edge aggregated graph attention layer, and finally 3) the node-level classification.

#### 2.2.1 Local feature extractor

A local feature extractor *λ* is applied, as shown in Figure 1(a), to a graph representation *G* of an arbitrary protein *P*. This layer is expected not only to capture the *local interactions* of the residues of the protein (sequence), but also to *reduce the dimension* of the node-level feature-vectors *q* = {*q*_1_, *q*_2_, *q*_3_, …, *q*_*N*_}. This helps the model learn to filter out unnecessary information, keeping the most useful information from *q*. Also, this helps the model to avoid overfitting, as this reduces the number of parameters for the subsequent layer.

**Figure 1:**
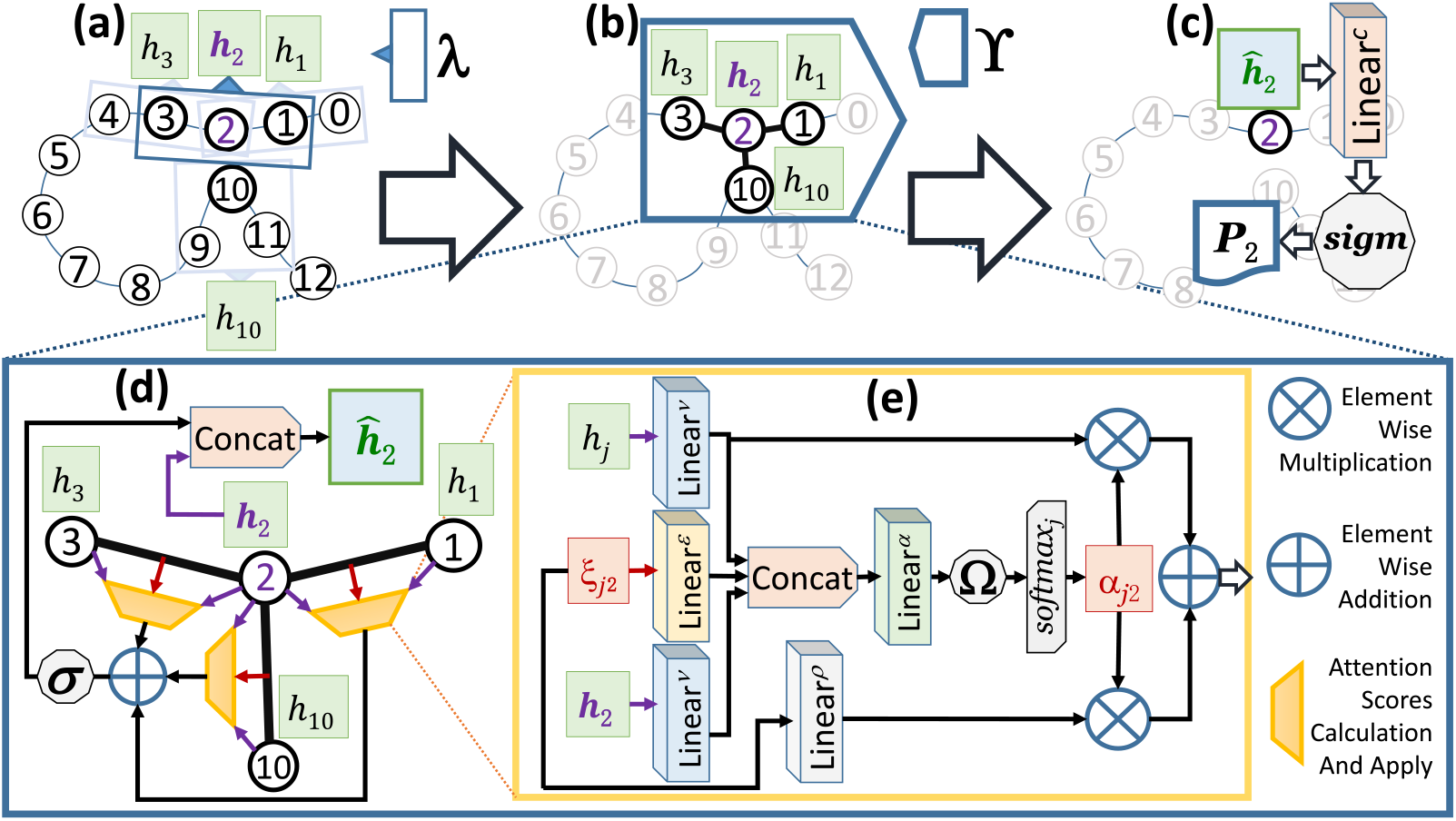
Schematic diagram of the overall pipeline of EGRET being applied to a dummy protein having 13 residues (*i* = 0, 1, 2, …, 12). (a) Local feature extractor *λ* (with window size *w*_*local*_ = 3) being applied. (b) Application of the edge aggregated graph attention layer ϒ to a neighborhood where residue 2 is the center node, and {1, 3, 10} ∈ 𝒩_2_ (neighborhood of node 2). (c) Node-level classifier applied on the final representation *ĥ*_2_ of node 2 (generated by ĥ). Here *sigm* represents the sigmoid activation function to generate a probability *P*_2_, which represents the numeric propensity of node 2 for being an interaction site. (d) The details on the edge aggregated graph attention layer ϒ shown in an expanded form (expansion is shown using dotted lines). Here, the yellow trapezoids represent the modules that calculate the attention scores and apply them to the features of the nodes and the edges. (e) The underlying working mechanism of a yellow trapezoid is shown in details in an expanded form. Here, *h*_*j*_ represents the feature representation of the node *j* ∈ 𝒩_2_, where *ξ*_*j*2_ represents the feature vector of the edge from node *j* to node 2. The *softmax*_*j*_ represents the softmax normalization applied to generate a normalized attention score *α*_*j*2_ for the edge from node *j* to node 2. In this figure, *σ* and Ω represent two activation functions, and **Linear**^*x*^ (*x* ∈ {*c, ν, ε, ρ, α*}) represents a linear layer with learnable parameter *W*^*x*^.

We used a one-dimensional convolutional neural network layer with a window size *w*_*local*_ as *λ*. Here, *w*_*local*_ is preferably a relatively small odd number to capture information about the relationship among the residues that are *sequentially closer*, but may or may not be closer in three-dimensional Euclidean space. The motivation behind taking an odd number is to ensure equal number of residues from both sides of a particular residue *i*. The sequence *q* of node features is passed through *λ* to generate a lower dimensional representation 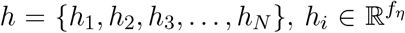, where *f*_*η*_ *< d*_*protbert*_. Here, for a residue *i, λ* encodes the feature-vectors {*q*_*j*_ |*q*_*j*_ ∈ *q and* 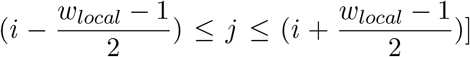} into a new condensed feature representation *h*_*i*_ for the node *i*.

#### 2.2.2 Edge aggregated graph attention layer

We now describe the original graph attention layer [52] and our proposed modifications by introducing edge aggregations. The feature representations *h* (generated by the local feature extractor, *λ*) are transformed using our proposed edge aggregated graph attention layer ϒ to encode the three-dimensional structural information of proteins.

In various Graph Neural Network based architectures [51, 52], there is an aggregation process where for a node *i* the feature representations of all the neighboring nodes N_*i*_ are aggregated to generate a fixed sized new representation *U*_*i*_ for the node *i*, which is then used for further computations. One highly used aggregation process is the weighted average of the features of the neighboring nodes. In such process, the representation of the *i*-th node 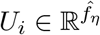 is generated according to the following Equation 1.

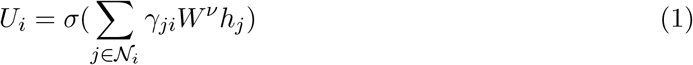

Here, 𝒩_*i*_ is the neighborhood of the node *i*. Also, *h*_*j*_ is the local feature representation of the node 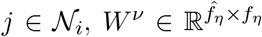 is a learnable parameter, and *γ*_*ji*_ is the weight that indicates how much important the features of the node *j* are to the node *i*. For Graph Convolutional Networks (GCN) [51] and Const-GAT [52], 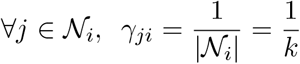, which is a constant. On the other hand, for GAT [52], *γ*_*ji*_ is a function ℱ(*·*) of the features *h*_*i*_ and *h*_*j*_ of the nodes *i* and *j*, respectively, representing the attention score of edge *E*_*ji*_ (see Eqn. 2). In 2018, Veličković *et al*. [52] showed the significant impact of these attention scores on the protein-protein interaction (PPI) dataset [60] consisting of graphs corresponding to different human tissues.

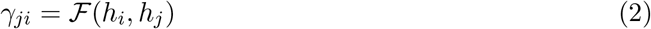

##### Using edge features during the calculation of the attention scores

In the calculation of the attention score as in Eqn. 2, in addition to node features 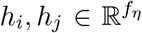, we incorporate *ξ*_*ji*_, the edge features of the directed edge from node *j* to node *i*. Equations 3 and 4 show the computations to generate the attention scores that are dependant not only on the nodes but also on the edges. Equation 3 represents a scoring function that is parameterized by a learnable parameter 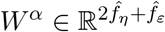, and Ω(*·*) is an activation function.

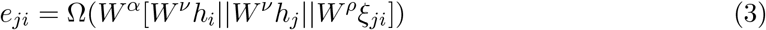

In Eqn. 3, *e*_*ji*_ is an unnormalized representation of the attention score and the symbol “||” represents the concatenation operation. Here, 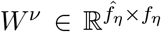 and 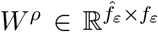 are learnable parameters used to apply linear transformation on the features of the nodes and the edges respectively.

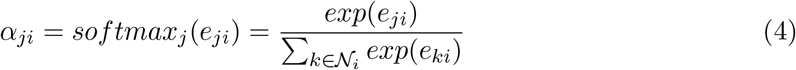

Equation 4 computes a softmax normalization (using the function *softmax_j*(*·*)) on {*e*_*ji*_|*j* ∈ 𝒩_*i*_} following [55]. This gives us a probability distribution over all the nodes *j* ∈ 𝒩_*i*_ (*i* being the center node) which is the attention distribution {*α*_*ji*_|*j* ∈ 𝒩_*i*_}.

##### Using edge features during the aggregation process

In order to utilize the full potential of the edge features *ξ*_*ji*_, we aggregate them alongside the features of the neighboring nodes {*h*_*j*_|*j* ∈ N_*i*_}, where node *i* is at the center of the neighborhood. We have updated Eqn. 1 accordingly and come up with Eqn. 5. Here, 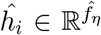 is the final feature representation of the node *i* after incorporating our proposed edge aggregated graph attention layer ϒ.

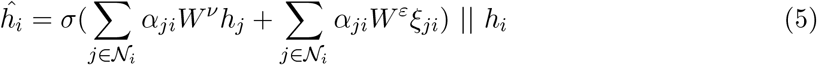

Here, *W*^*ν*^ is the same learnable parameter as in Equation 3, and 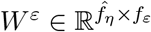 is a new learnable parameter, which is multiplied with the edge feature vector *ξ*_*ji*_ to apply a linear transformation on *ξ*_*ji*_ before aggregation. Here, *σ*(*·*) is an activation function and the output of *σ*(·) is concatenated with *h*_*i*_ (the previous feature representation of the node *i*) and thereby generating the new representation *ĥ*_*i*_, which is the output of our edge aggregated graph attention layer ϒ. Figure 1(b) demonstrates ϒ being applied on a neighborhood of a dummy protein. Figures 1(d) and 1(e) show the detailed mechanism of ϒ layer.

#### 2.2.3 Node-level classification

For each node *i* ∈ *V* (*G*), the final feature representation *ĥ*_*i*_ (generated by edge aggregated graph attention layer ϒ) is linearly transformed to produce a single scalar followed by an application of the sigmoid activation function as shown in Eqn. 6. This generates a probability 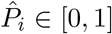, representing the numeric propensity 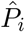 of residue *i* for interactions with other proteins. Here, 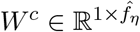 is a learnable parameter and *sigm*(·) is the Sigmoid activation function [61]. Figure 1(c) shows the node-level classification in EGRET.

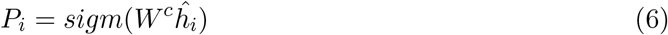

### 2.3 Overall end-to-end pipeline of EGRET

Figure 1 shows the overall end-to-end pipeline of EGRET. Here we demonstrate EGRET being applied on a dummy protein with thirteen residues and show the computation for only residue 2 for the sake of readablity and clarity of this figure. EGRET starts with representing a protein as a graph *G*, where each node *i* ∈ *V* (*G*) is connected to |𝒩_*i*_| = *k* other closest nodes with directed edges. For the sake of readability, we used |𝒩_*i*_| = *k* = 3 in this example. Here 𝒩_2_ = {1, 3, 10}. Note that residue 10 is not sequentially closer to node 2, but is in close proximity of node 2 in three-dimensional space.

EGRET converts the residues to a series of tokens (each token representing a residue) and uses ProtBERT [43] to generate an embedding-vector for each of the residues. These embedding-vectors are assigned as the initial feature-representations *q* of the nodes in *V* (*G*). Alongside, the edge feature vectors {*ξ*_*j,i*_|*j, i in V* (*G*)} are calculated from the structural data of the protein (available in PDB files [50]). Each feature-vector *ξ*_*j,i*_ is associated with one directed edge *E*_*j,I*_ from node *j* to node *i*. Next, local feature extractor *λ* is applied to the feature-representations (*q*) of the nodes of the proteins. *λ* generates a new feature representation *h*_*i*_ of a residue *i* (see Figure 1(a)). The details of local feature extraction have been described in Sec. 2.2.1.

Once the local feature extraction is completed, the edge aggregated graph attention layer ϒ is applied on each of the neighborhoods. We show the application of ϒ only to the neighborhood of residue 2 in Fig. 1(b)). Please see Sec. 2.2.2 for details. ϒ generates the final feature representation *ĥ*_*i*_ of the central node *i* of a neighborhood. Finally, node-level classification (Sec. 2.2.3) is applied to the final representation (*ĥ*_2_ in this example). Node level classifier provides us with a probability value *P*_2_ for residue 2, which is the predicted propensity of residue 2 being an interfacing residue (or interaction site). This same end-to-end pipeline is applied to all other residues and thereby computing the propensities of all the residues.

### 2.4 GAT-PPI: GAT based PPI site prediction without edge aggregation

As this is the first known study on leveraging graph neural networks for PPI site prediction for single proteins, we have developed an original GAT [52] based PPI site prediction approach without any edge aggregation in order to show the superiority of graph based approach over other competing approaches. We call this method GAT-PPI. We used the original implementation provided by the Deep Graph Library [62].

## 3 Results and Discussion

### 3.1 Dataset

We evaluated EGRET on three previously assembled benchmark datasets. The first dataset (henceforth referred to as the DeepPPISP dataset) is a combination of three widely used benchmark datasets, namely (1) Dset_186 [29], (2) Dset_72 [29], and (3) PDBset_164 [28]. Dset_186 and Dset_72 [29], contain 186 (from 105 heterodimeric protein complexes) and 72 non-repetitive protein sequences (from 36 protein complexes in the protein–protein docking benchmark set version 3.0 [63]), respectively. PDBset_164 [28] includes 164 protein sequences (all of them are heterodimers with sequence length ≥ 50). In these datasets, a residue is considered a *surface residue* if its relative solvent accessibility (rSA) is at least 5% [33], and a surface residue is defined as an *interaction site* if its reduction of the absolute solvent accessibility is at least 1 Å^2^ on complex formation [33] (see Table 1 for the distribution of interaction and non-interaction sites in these datasets).

**Table 1:** summary of the deepppisp dataset analyzed in this study.

| Dataset | Proteins | Interaction sites | Non-interaction sites |
| --- | --- | --- | --- |
| Dset_186 | 186 | 5,517 (15.23%) | 30,702 (84.77%) |
| Dset_72 | 72 | 1,923 (10.60%) | 16,217 (89.40%) |
| PDBset_164 | 164 | 6,096 (18.10%) | 27,585 (81.90%) |
| Train+Validate | 352 | 11,079 (15.14%) | 62,102 (84.86%) |
| Test | 70 | 2,332 (19.78%) | 9,459 (80.22%) |

All these three datasets have been built with proteins from PDB-database [50], with sequence similarity less than 25% and resolution less than 3.0 Å (solved by X-ray crystallography). We refer to [29] and [28] for more details. As these datasets come from different research groups, the authors of DeepPPISP [15] integrated the three datasets into a fused dataset to ensure that the training set and the test set are from an identical distribution. Zeng *et al*. [15] split this fused dataset into a test set comprising 70 randomly selected protein sequences and a training set with the remaining (about 83.4%) protein sequences. The training set contains 352 protein sequences, 50 of which are used for independent validation. They evaluated their method DeepPPISP as well as other state-of-the-art methods on this split using the same evaluation scheme. For the sake of a fair comparison, we used the same splitting and evaluation scheme used by [15]. Thus, EGRET was trained on 302 proteins in the training set, followed by a validation on the 50 hold-out proteins to tune the hyper parameters. Finally, it was tested on the test set comprising 70 proteins. Table 1 shows the numbers of interaction and noninteraction sites in the DeepPPISP datasets and the splits used in this study [15]. We note that DELPHI [25], which is a sequence-based method, used a much larger dataset containing 9,982 protein sequences for training and validation. However, leveraging that large dataset for training structure-based methods is difficult due to the unavailability of curated structural information.

In addition to the DeepPPISP benchmark dataset, we also evaluated EGRET on a substantially larger dataset compiled by the authors of MaSIF [24]. This dataset was assembled from the PRISM dataset [64, 65], the ZDock benchmark [66], PDBBind [67] and the SAbDab antibody database [68]. MaSIF dataset contains 3362 proteins, filtered according to sequence identity using the psi-cd-hit60 [69] at 30% sequence identity. Finally, we assessed the performance of EGRET on the Dockground [70] dataset which is a benchmark dataset for evaluating protein-protein docking methods. This is a comprehensive resource including co-crystallized (bound) protein complexes, unbound X-ray and simulated docking benchmark sets, model–model docking benchmark sets, scoring benchmarks (docking decoys), and templates for comparative docking.

### 3.2 Methods compared

We compared our proposed EGRET and GAT-PPI (the proposed model without edge aggregation) with state-of-the-art sequence-based as well as structure-based methods. We considered nine competing methods for predicting PPI sites, namely SPPIDER [21], ISIS [36], PSIVER [29], SPRINGS [28], RF_PPI [18], and especially the most recent and the most accurate predictors IntPred [16], SCRIBER [26], DeepPPISP [15] and DELPHI [25]. Among these nine alternate methods, SPIDER, IntPred and DeepPPISP are strucuture-based methods that leverage structural data or features derived from structural data. See supplementary materials for further details. We also compared EGRET with MaSIF and PInet [20] on the MaSIF benchmark dataset.

### 3.3 Evaluation metrics

For the evaluation of EGRET, we used seven widely used evaluation metrics [15, 25, 26], namely accuracy, precision, recall, F1-measure (F1), area under the receiver operating characteristic curve (AUROC), area under the precision-recall curve (AUPRC), Matthews correlation coefficient (MCC). See supplementary materials for more details.

Note that accuracy is not a meaningful evaluation metric for unbalanced dataset. As rightly mentioned by [25], AUROC and AUPRC convey a comprehensive performance measurement of a method since these two metrics are threshold independent. Among the other metrics, F1-measure and MCC are the most important performance metrics since PPISP is an imbalanced learning problem [15, 71, 72]. We performed Wilcoxon signed-rank test [73] (with *α* = 0.05) to measure the statistical significance of the differences between two methods.

### 3.4 Results on DeepPPISP benchmark dataset

The comparison of EGRET with other state-of-the-art methods are shown in Table 2. Remarkably, EGRET outperformed all other methods under six (out of seven) evaluation metrics, including the most important ones (e.g., F1, AUROC, AUPRC, MCC). Recall is the only metric where EGRET was outperformed by GAT-PPI (which is our proposed model without edge aggregation). Notably, GAT-PPI also achieved the second best performance on the four important evaluation metrics, namely F1, AUROC, AUPRC, MCC, which are second to only EGRET. Thus, our proposed GAT based models EGRET (with edge aggregation) and GAT-PPI (without edge aggregation) are the best and second-best methods, respectively on this benchmark dataset. These results clearly show the superiority of the proposed GAT based architecture (with or without edge aggregation) over the best alternate methods. The F1-score as well as AUPRC and MCC – two of the most important evaluation metrics due to the imbalance nature of PPI prediction problem – obtained by EGRET are 0.438, 0.405 and 0.27, respectively, which are 4.8%, 12.5% and 14.4% higher than those achieved by the best existing method DELPHI, and these improvements are statistically significant (*p*-value *<* 0.05). See Table S2 in supplementary materials.

**Table 2:** A comparison of the predictive performance of our proposed EGRET and GAT-PPI with other state-of-the-art methods on the DeepPPISP benchmark dataset. The best and the second best results for each metric are shown in bold and italic, respectively. Values which were not reported by the corresponding source are indicated by “-”.

| Method | ACC | Precision | Recall | F1 | AUROC | AUPRC | MCC |
| --- | --- | --- | --- | --- | --- | --- | --- |
| SPPIDER<sup>1,2</sup> | 0.622 | 0.209 | 0.459 | 0.287 | - | 0.23 | 0.089 |
| ISIS<sup>2</sup> | *0.694* | 0.211 | 0.362 | 0.267 | - | 0.24 | 0.097 |
| PSIVER<sup>2</sup> | 0.653 | 0.253 | 0.468 | 0.328 | - | 0.25 | 0.138 |
| SPRINGS<sup>2</sup> | 0.631 | 0.248 | 0.598 | 0.35 | - | 0.28 | 0.181 |
| RF_PPI<sup>2</sup> | 0.598 | 0.173 | 0.512 | 0.258 | - | 0.21 | 0.118 |
| IntPred<sup>1,2</sup> | 0.672 | 0.247 | 0.508 | 0.332 | - | - | 0.165 |
| SCRIBER | 0.616 | 0.274 | 0.569 | 0.37 | 0.635 | 0.307 | 0.159 |
| DeepPPISP<sup>1,2</sup> | 0.655 | 0.303 | 0.577 | 0.397 | 0.671 | 0.32 | 0.206 |
| DELPHI | 0.667 | *0.32* | *0.604* | 0.418 | 0.69 | 0.36 | 0.236 |
| GAT-PPI<sup>1</sup> | 0.653 | 0.318 | **0.659** | *0.429* | *0.714* | *0.398* | *0.252* |
| EGRET<sup>1</sup> | **0.715** | **0.358** | 0.561 | **0.438** | **0.719** | **0.405** | **0.27** |

<sup>1</sup> Uses structural information.
<sup>2</sup> Results reported by DeepPPISP [15].

Notably, our proposed GAT based model GAT-PPI, even without the edge aggregation, outperformed other existing methods including DELPHI and DeepPPISP, and the improvements are statistically significant. EGRET is substantially better than two of the most recent structure-based methods, namely DeepPPISP and IntPred. EGRET achieved 10.3%, 7.2%, 26.6% and 31.1% higher scores for the above-mentioned metrics respectively than those of DeepPPISP. In particular, 26.6% and 31.1% improvement over DeepPPISP in two of the most important metrics AUPRC and MCC is quite remarkable. Note that ISIS, which achieved the second highest accuracy, had the lowest F1 measure and second lowest MCC, showing that accuracy is not a meaningful evaluation metric for unbalanced datasets.

In order to visually show the predicted interfacial sites, we show in Fig. 2 the true and predicted (by EGRET and DELPHI) interaction sites on a representative protein (PDB ID 3OUR, Chain-B). This protein is 150 residue long with 39 interaction sites. It has 412 non-local contact pairs, suggesting a high level of long-range interactions.

**Figure 2:**
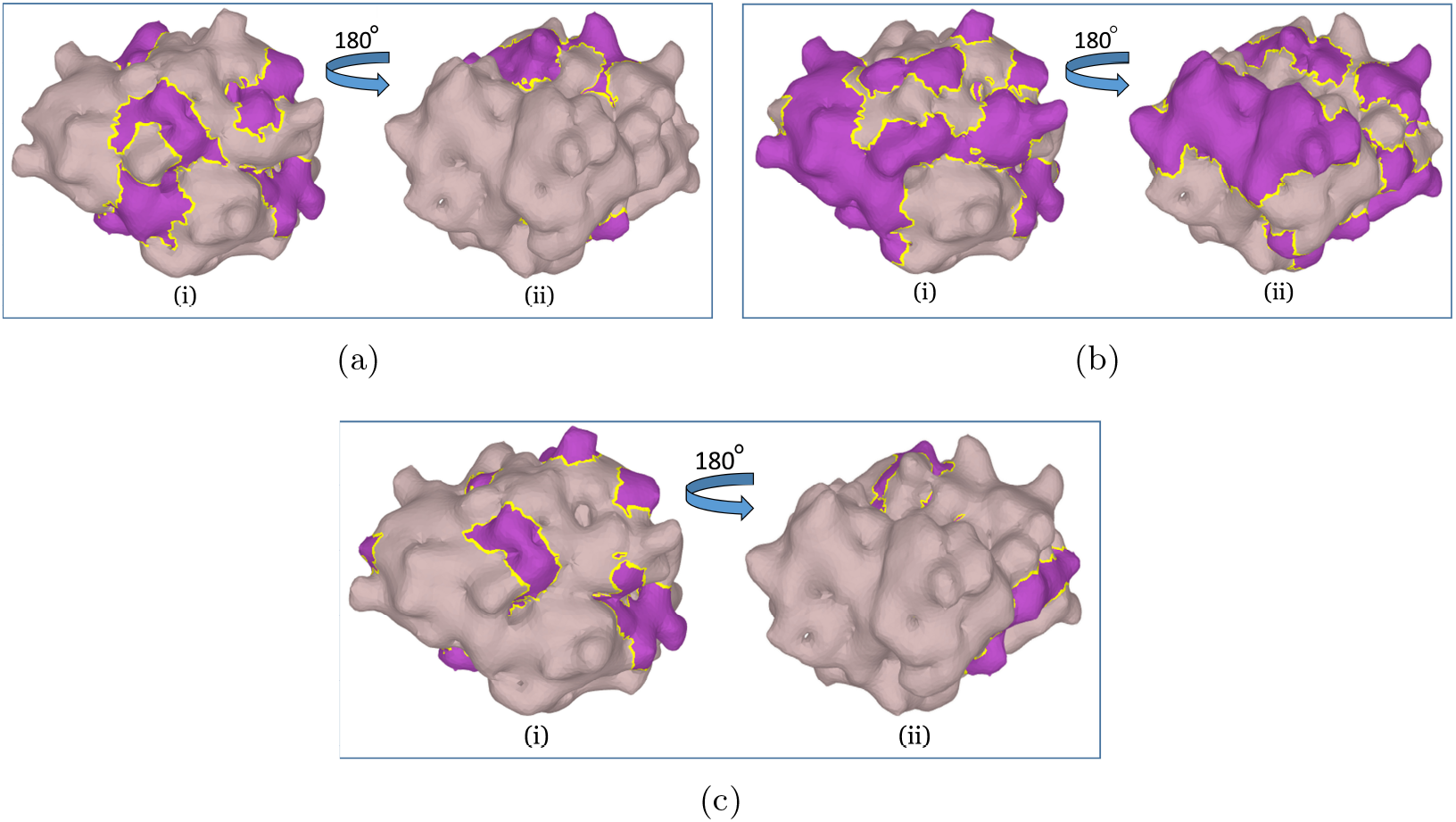
Interaction sites of a representative protein in the test set (PDB ID 3OUR, Chain-B). (a) Interaction sites predicted by EGRET, (b) interaction sites predicted by DELPHI, and (c) true interactions sites as obtained from the dataset. Interaction sites are shown in purple on the protein surface. The left and right images (i and ii) in each of the figures (a, b and c) show two opposite sides (i.e., 180° rotated view).

### 3.5 Impact of long-range interactions in PPI sites prediction

EGRET, unlike DeepPPISP, is designed for generating different suitable global features for different residues. Therefore, we investigated the performance of EGRET under various levels of long-range interactions, and compared with Delphi and GAT-PPI. We could not include DeepPPISP in this experiments as its protein-wise predictions are not publicly available and the webserver at http://bioinformatics.csu.edu.cn/PPISP/ is not accessible (last accessed on Oct 10, 2020). The predictions of DELPHI were obtained from DELPHI webserver (available at: https://delphi.csd.uwo.ca/).

We computed the number of non-local interactions per residue for each of the 70 proteins in our test set, and sorted them in an ascending order. Two residues at sequence position *i* and *j* are considered to have non-local interactions if they are at least 20 residues apart (|*i* − *j*| ≥ 20), but *<* 8 *Å* away in terms of their atomic distances between *Cα* atoms [74]. Next, we put them in seven equal sized bins *b*_1_, *b*_2_, …, *b*_7_ (each containing 10 proteins, where *b*_1_ contains the proteins with the lowest level of non-local interactions (0.41-1.49 non-local contact per residue) and *b*_7_ represents the model condition with the highest level of non-local interactions (2.59-3.21 non-local contact per residue). We show the AUPRC obtained by EGRET, GAT-PPI and DELPHI under various model conditions in Fig. 3 (a) and Tables S3 and S4 in supplementary materials. These results show that – as expected – the performance of GAT-PPI, EGRET and DELPHI degrades as we increase the number of non-local contacts. However, the difference in predictive performance between GAT-PPI (or EGRET) and DELPHI significantly increases with increasing levels of non-local interactions (with a few exceptions). Notably, there is no statistically significant difference between EGRET (or GAT-PPI) and DELPHI on *b*_1_ (*p >* 0.05), but as we increase the level of non-local interactions, EGRET and GAT-PPI tend to become more accurate than DELPHI and attain significant improvement (*p <* 0.05) on *b*_7_ (see Supplementary Tables S3 and S4). These results indicate that addressing non-local interactions by suitable global features is one of the key factors in the improvement.

**Figure 3:**
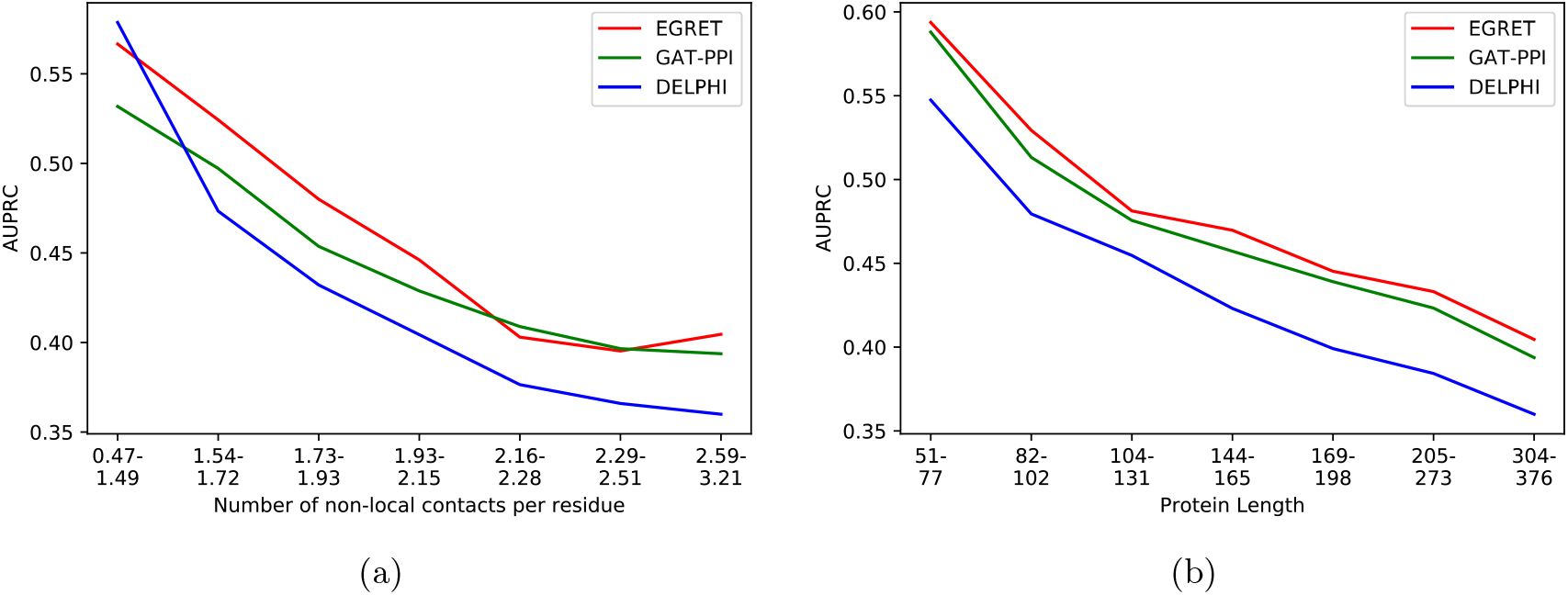
Impact of long-range interactions and protein lengths on predictive performance of PPI sites prediction. (a) AUPRC of EGRET, GAT-PPI and DELPHI on varying levels of non-local interactions. (b) AUPRC of EGRET, GAT-PPI and DELPHI on varying lengths of the proteins.

### 3.6 Impact of protein length

We investigated the impact of protein lengths since we took global features into consideration. We divided 70 proteins in our test set into seven non-overlapping bins based on their lengths. We observed a similar trend as in long-range interactions – the predictive performance deteriorates with increasing lengths of the proteins (see Fig. 3 (b)) and Tables S5 and S6 in supplementary materials. EGRET and GAT-PPI consistently outperform DELPHI across various model conditions and the improvement tend to increase (with a few exceptions) and become statistically significant as we increase the length of the proteins.

### 3.7 Impact of edge aggregation in graph attention network

The initial enthusiasm for using edge features was to assist the network in generating an embedding with *richer structural information* for each of the nodes in the graph. Indeed, edge aggregation has a significant impact on PPI sites prediction as supported by the experimental results shown in Table 2. EGRET achieved better performance metrics than GAT-PPI (except for the recall). It obtained 9.5%, 12.6%, 2.1%, 0.7%, 1.8%, 7.1% performance improvement over GAT-PPI in accuracy, Precision, F1-score, AUROC, AUPRC, and MCC, respectively. These improvements (albeit small for some of the evaluation metrics) indicate the positive impact of edge aggregation.

### 3.8 Impact of transfer learning using ProtBERT-based features

In order to show the impact of transfer learning, we investigated the efficacy of the embeddings of the nodes (residues) generated by ProtBERT compared to other types of feature representations that have been widely used in the PPI literature. We compared the impact of DeepPPISP Features containing PSSM, raw sequence features, and eight-state secondary structure features with ProtBERT-based embeddings. We trained EGRET and GAT-PPI using both these feature sets (ProtBERT-based features and DeepPPISP features) and analyzed their predictive performance (see Table 3). The results suggest that both GAT-PPI and EGRET obtained better perfomance with the ProtBERT-based features than those achieved with DeepPPISP features, indicating positive effects of transfer learning. Notably, EGRET consistently outperformed GAT-PPI on both feature sets – suggesting the positive impact of edge aggregation regardless of the choice of feature sets. Moreover, even without the ProtTrans features (i.e., with the DeepPPISP feature set), EGRET is better than or as good as DeepPPISP and DELPHI (See Tables 2 and 3). Informatively, EGRET used only 352 proteins for training and validation, whereas DELPHI used a substantially larger dataset with 9,982 proteins. Therefore, matching or improving upon the accuracy of DELPHI without transfer learning is noteworthy.

**Table 3:**
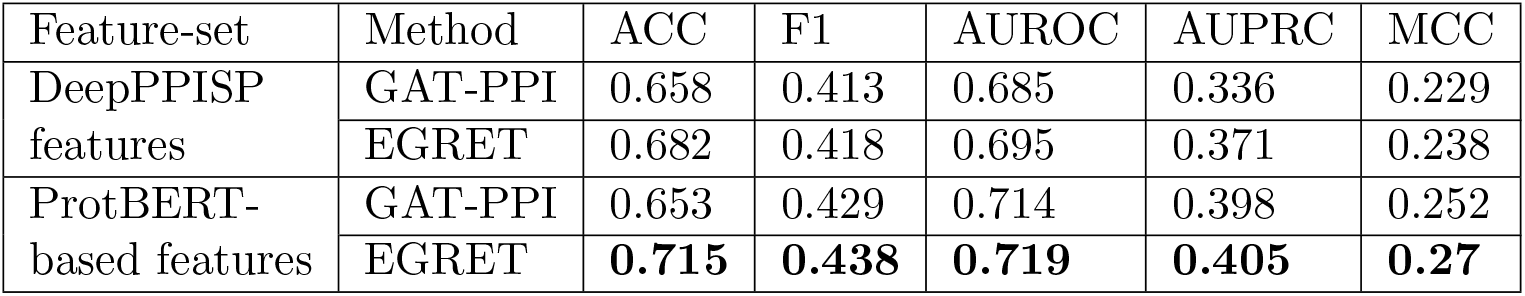
Impact of different types of features. We show the performance of EGRET and GAT-PPI on DeepPPISP feature set and ProtBERT-based feature set. Best results are shown in bold.

| Feature-set | Method | ACC | F1 | AUROC | AUPRC | MCC |
| --- | --- | --- | --- | --- | --- | --- |
| DeepPPISP features | GAT-PPI | 0.658 | 0.413 | 0.685 | 0.336 | 0.229 |
|  | EGRET | 0.682 | 0.418 | 0.695 | 0.371 | 0.238 |
| ProtBERT-based features | GAT-PPI | 0.653 | 0.429 | 0.714 | 0.398 | 0.252 |
|  | EGRET | **0.715** | **0.438** | **0.719** | **0.405** | **0.27** |

### 3.9 Evaluation on the MaSIF dataset

The authors of MaSIF used 3003 proteins for training and 359 for testing. Later, the authors of PInet [20] removed the complexes whose interaction sites was *<* 1% of the size of the ligand, resulting in a training-test split of 2689 and 345 proteins. We compared EGRET with MaSIF and PInet with the same training-test split. We note that PInet is a pairwise PPISP prediction method in contrast to MaSIF and EGRET that are partner-independent method for predicting putative interaction sites upon the surface of an isolated protein.

The authors of MaSIF and PInet trained their models with the following two types of features: i) geometric features only (i.e., structural features) and ii) geometric features and physicochemical properties (e.g., electrostatics and hydrophobicity). We compared EGRET to MaSIF and PInet trained with both types of features. The results are presented in Table 4 which indicate that EGRET achieved higher AUPRC than PInet (with geometric features alone), but was worse than PInet with full set of features. In terms of AUROC, PInet and MaSIF outperformed EGRET with the full set of features, but EGRET outperformed both PInet and MaSIF when only geometric features were used. We note that AUPRC is more meaningful than AUROC for such imbalanced datasets. We further note that PInet, being a pairwise PPISP method, makes use of information from the partner proteins. Therefore, a direct comparison with PInet is inappropriate. Remarkably, however, EGRET outperforms PInet trained on geometric features alone.

**Table 4:**
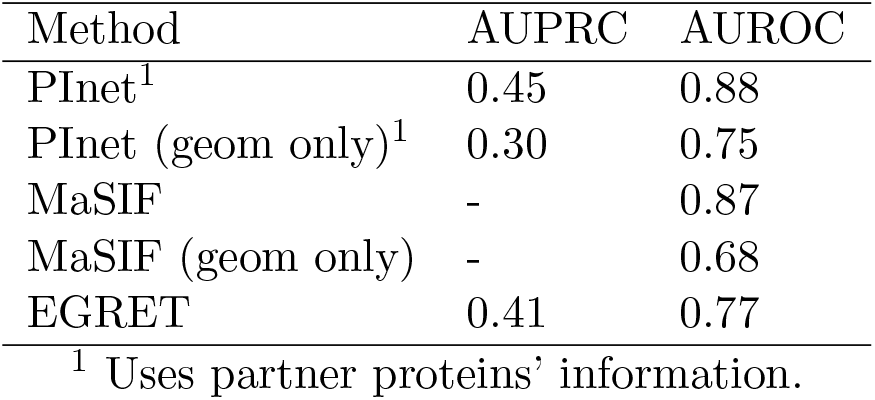
Performance in terms of the area under PR and ROC curves on the MaSIF benchmark dataset. Results of PInet and MaSIF were obtained from [20]. AUPRC of MaSIF was not reported by the corresponding source.

| Method | AUPRC | AUROC |
| --- | --- | --- |
| PInet<sup>1</sup> | 0.45 | 0.88 |
| PInet (geom only)<sup>1</sup> | 0.30 | 0.75 |
| MaSIF | - | 0.87 |
| MaSIF (geom only) | - | 0.68 |
| EGRET | 0.41 | 0.77 |

<sup>1</sup> Uses partner proteins' information.

### 3.10 Evaluation on the Dockground dataset

Dockground [70] is a benchmark dataset for evaluating protein-protein docking methods. Although EGRET is partner-independent, predicting putative binding sites in general for an isolated protein, rather than making predictions that are specific to given partner proteins, evaluating its performance on the Dockground dataset will help us investigate if the interaction sites corresponding to different complexes as obtained from the Dockground dataset are contained within the likely interaction sites predicted by EGRET. We take the union of the sets of interaction sites corresponding to various complexes of a particular protein in the Dockground dataset, except for the interacting residues within the biounit chains of that protein (when a protein has more than one chains). For example, consider the protein 1GY6_AB, for which we want to find the interaction sites from the Dockground dataset. 1GY6_AB is present in the complex 1A2K as biounit chains A and B. In the Dockground dataset, there are labeled interactions sites for five chains A, B, C, D, E in 1A2K. While finding the set of the interaction sites of 1GY6_AB in the complex 1A2K, we consider the pairs (A, C), (A, D), (A, E), (B, C), (B, D), (B, E), but not (A, B). The interaction sites thus obtained are compared to those predicted by EGRET. In this case, recall is a more meaningful evaluation metric than precision. We evaluated EGRET on 25 proteins in this dataset (see Table 5). The recall of EGRET was greater than 0.9 in the majority of cases, and in some cases it was equal to or close to one. The other measures produced by EGRET (e.g., F1, AUROC, AUPRC, accuracy) were also reasonably high, and even higher than those obtained on the DeepPPISP benchmark data (i.e., Dset_186 [29], Dset_72 [29], and PDBset_164 [28]). For example, the F1 score and precision of EGRET on the DeepPPISP benchmark data were 0.438 and 0.358, respectively (see Table 2), whereas they are substantially higher on the proteins in the Dockground dataset. These results suggest that EGRET can predict a useful set of putative interaction sites which can be considered as a preamble of the more elaborated docking method.

**Table 5:** Performance of EGRET on 25 representative proteins from the Dockground dataset [70]. Here the “Label sources in Dockground” column shows which complexes were used to find the set of interacting residues in the Dockground dataset.

| Protein Name<br>(Format:<br>[Complex’s PDB<br>ID]-[Biounit<br>Chain Names]) | Label sources in Dock-<br>ground<br>(Format: [Complex’s<br>PDB ID]) | F1 | Precision | Recall | AUROC | AUPRC |
| --- | --- | --- | --- | --- | --- | --- |
| 1GY6_AB | 1A2K | 0.450 | 0.290 | 1.0 | 0.486 | 0.256 |
| 3RAN_A | 1A2K | 0.642 | 0.515 | 0.850 | 0.938 | 0.486 |
| 1I7Y_AB | 1I7Y | 0.628 | 0.509 | 0.823 | 0.758 | 0.610 |
| 1BKF_A | 1B6C, 3FAP, 1BKF | 0.776 | 0.691 | 0.884 | 0.863 | 0.746 |
| 1DOK_A | 1DOK, 4DN4, 1ML0 | 0.796 | 0.740 | 0.860 | 0.736 | 0.781 |
| 1DDJ_A | 1BUI, 1BUI | 0.553 | 0.483 | 0.646 | 0.717 | 0.507 |
| 2SAK_A | 1BUI | 0.554 | 0.387 | 0.979 | 0.407 | 0.341 |
| 1IXM_AB | 1F51 | 0.399 | 0.249 | 1.0 | 0.410 | 0.199 |
| 1MH1_A | 1E96, 1G4U, 1HE1,<br>1I4D, 2H7V, 2NZ8 | 0.564 | 0.405 | 0.927 | 0.746 | 0.649 |
| 1OGW_A | 1P3Q, 1S1Q, 1UZX,<br>1WR6, 1WRD, 2AYO | 0.881 | 0.857 | 0.906 | 0.894 | 0.961 |
| 3LZT_A | 1A2Y, 1GPQ, 1MLC,<br>1NBY, 1SQ2, 1UUZ | 0.810 | 0.750 | 0.880 | 0.789 | 0.820 |
| 2CE2_X | 1HE8, 1K8R, 1LFD,<br>1NVU, 1WQ1 | 0.713 | 0.571 | 0.947 | 0.780 | 0.717 |
| 2FXU_A | 1MA9, 1Y64, 2A41,<br>2A42, 2PAV | 0.518 | 0.356 | 0.950 | 0.608 | 0.420 |
| 1C3D_A | 1GHQ, 2WY7, 3OED,<br>3RJ3 | 0.523 | 0.423 | 0.682 | 0.700 | 0.475 |
| 1GZ8_A | 1BUH, 1F5Q, 1OIU,<br>1W98 | 0.449 | 0.647 | 0.344 | 0.702 | 0.526 |
| 1P9A_G | 1OOK, 1P8V, 1SQ0,<br>1U0N | 0.555 | 0.424 | 0.803 | 0.657 | 0.363 |
| 1QG2_A | 1I2M, 1IBR, 1K5D,<br>1QBK | 0.326 | 0.316 | 0.336 | 0.211 | 0.377 |
| 1YJ1_A | 1NBF, 1XD3, 2G45,<br>2OOB | 0.745 | 0.792 | 0.704 | 0.842 | 0.810 |
| 2OVO_A | 1CHO, 1HJA, 1PPF,<br>2SGE | 0.711 | 0.727 | 0.696 | 0.769 | 0.703 |
| 2FAD_A | 4KEH, 4ETW, 4IHF | 0.750 | 0.618 | 0.955 | 0.889 | 0.659 |
| 1D3Z_A | 3PRP, 4WLR, 4BOZ | 0.783 | 0.964 | 0.659 | 0.817 | 0.886 |
| 2XQW_A | 2XQW, 3D5R, 2WY8 | 0.384 | 0.238 | 1.0 | 0.579 | 0.284 |
| 1U9B_A | 1Z5S, 2O25, 1KPS | 0.762 | 0.619 | 0.990 | 0.566 | 0.648 |
| 1KMQ_A | 4XOI, 1TX4, 1CXZ | 0.671 | 0.533 | 0.903 | 0.816 | 0.711 |
| 2GKR_I | 1SGP, 1R0R, 1PPF | 0.651 | 0.609 | 0.700 | 0.634 | 0.549 |

We found twelve proteins that are common in both the Dockground dataset and the DeepPPISP benchmark dataset. These DeepPPISP benchmark datasets are relatively old and the PDB content has grown with new protein-protein complexes. The number of interaction sites in the Dockground dataset has increased for two of these 12 proteins. We investigated the performance of EGRET on these two proteins. EGRET performed substantially better with interaction sites obtained from the Dockground dataset compared to those obtained from the DeepPPISP benchmark data (see Table 6). Many of the predicted interaction sites that were false positives in DeepPPISP data turned out to be true positives in Dockground data. The average precision, F1-score, and AUPRC of EGRET improved from 0.506 to 0.716, 0.634 to 0.786, and 0.515 to 0.764, respectively, with new set of interaction sites in the Dockground data. It is worth noting that the known interaction sites represent a fraction of all possible putative interaction sites. With the discovery of new protein-protein interactions, the set of known interaction sites for a specific protein may grow. As such, the improved precision of EGRET on the Dockground dataset (when compared to the relatively old DeepPPISP data) suggests that the set of putative interaction sites predicted by EGRET are indeed very likely to interact with other proteins.

**Table 6:** Performance of EGRET on two proteins that are common in the DeepPPISP test set and the Dockground dataset, but with different numbers of labeled interacting residues. Performance metrics were computed with respect to the interaction sites from the DeepPPISP dataset and the Dockground dataset. Here the “Label sources” column shows which complexes were considered to find the set of interacting residues.

| Dataset considered for labeling | Protein Name<br>(Format:<br>[Complex’s<br>PDB<br>ID]_[Biounit<br>Chain<br>Names]) | Label sources<br>(Format:<br>[Complex’s<br>PDB ID]) | F1 | Precision | Recall | AUROC | AUPRC |
| --- | --- | --- | --- | --- | --- | --- | --- |
| DeepPPISP dataset | 1B6C_A | 1B6C | 0.656 | 0.556 | 0.800 | 0.819 | 0.454 |
|  | 1ML0_D | 1ML0 | 0.612 | 0.455 | 0.938 | 0.813 | 0.575 |
| Dockground dataset | 1B6C_A | 1B6C, 3FAP, 1BKF | 0.776 | 0.691 | 0.884 | 0.863 | 0.746 |
|  | 1ML0_D | 1ML0, 1DOK, 4DN4 | 0.796 | 0.740 | 0.860 | 0.736 | 0.781 |

### 3.11 Consistency of the EGRET architecture

We investigated the behavior of EGRET to provide insights on how the architecture is making decisions. Some recent studies demonstrated the behavior and partial interpretability of deep neural networks for solving various problems in computational biology (in particular, see [40] and [48]).

#### 3.11.1 Correlation between the neighboring nodes (consistency of the local representations)

Let the EGRET predicted numeric propensity of interaction for any residue *r* be *P*_*r*_ ∈ [0, 1], and the true and predicted labels of *r* be *Y*_*r*_ and *Ŷ*_*r*_, respectively. The joint predicted probability of two residues *i* and *j* is *P*_*i*_ \**P*_*j*_. Let 𝒢_*prot*_ = {*G*_1_, *G*_2_, …, *G*_*N*_} be the set of *N* graphs representing *N* proteins in our test set, and *V* (𝒢_*prot*_) and *E*(𝒢_*prot*_) represent the sets of nodes and edges in 𝒢_*prot*_, respectively. In the following analyses, (*a*_*i*_), *i* ∈ *V* (𝒢_*prot*_), represents an ordered list (sequence) that is ordered by the value of *i*, where *a*_*i*_ may represent *Y*_*i*_, *Ŷ*_*i*_ or *P*_*i*_.

We investigated the correlation between numeric propensities corresponding to the source and destination nodes of the directed edges in our proposed graph based model. More specifically, for an edge {*E*_*ij*_|*E*_*ij*_ ∈ *E*(𝒢_*prot*_) *and i, j* ∈ *V* (𝒢_*prot*_)}, we computed the correlation coefficient between the two ordered lists (*P*_*i*_) and (*P*_*j*_). We ran this analysis on the entire test set of the DeepPPISP benchmark data using the Pearson correlation function. We found that there is a high positive correlation between them (correlation coefficient *r* = 0.782), and this correlation is statistically significant with *p*-value ≪ 0.05 (according to the Spearman rank-order correlation). This shows us that two nodes are likely to be predicted as the same type (either interaction or non-interaction site) by EGRET, if there is an edge between them. This demonstrated correlation between numeric propensities corresponding to the source and destination nodes of an edge is largely due to the fact that interaction and non-interaction sites usually belong to contiguous regions in the protein structure. As such, there is a meaningful relationship between the model behavior/architecture and biological properties of the interaction sites.

#### 3.11.2 Patterns of attention scores

In order to investigate the impact of edges and their weights (i.e., attention scores) and how effectively the network learns to share information, we investigate – for each residue *i* ∈ *V* (𝒢_*prot*_) which is predicted to be an interaction site by EGRET – how much the predictions of its neighbors correlate with their corresponding edge weights (i.e. the attention scores). More specifically, ∀_*i*_{*i* ∈ *V* (𝒢_*prot*_) *and Ŷ*_*i*_ = 1} we compute the correlation coefficient between the predictions of the neighbors 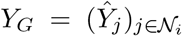 and the associated attention scores *A*_*G*_ = (*α*_*ji*_). We found a positive correlation coefficient (*r* = 0.243) between *Ŷ*_*G*_ and *A*_*G*_, which is statistically significant with *p*-value ≪ 0.05 (according to the Spearman rank-order correlation). This suggests that for a particular interaction site, its neighbors with relatively higher attention scores are more likely to be an interaction site than its neighbors with relatively lower attention scores. We further demonstrate this correlation with a cartoon figure using a representative protein (PDB ID 3OUR, Chain-B), available in the test set (see Fig. 4(a)).

**Figure 4:**
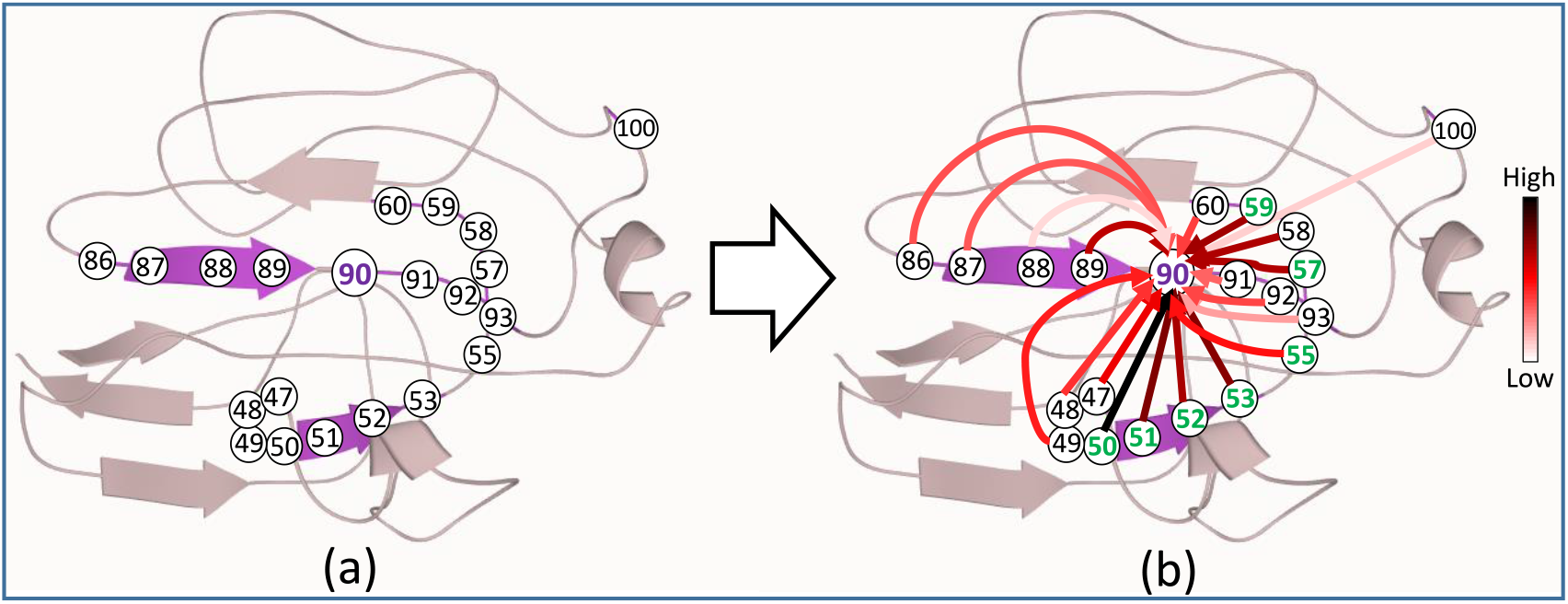
Patterns of attention scores produced by EGRET in predicting the interaction sites. (a) A cartoon images of a protein (PDB ID 3OUR, Chain B), where we show the neighborhood (window size = 20) of the residue 90. (b) The attention scores along the edges in this neighborhood using a color gradient which vary continuously from light red to black with increasing attention scores. The residues shown by green nodes are the interaction sites predicted by EGRET.

Figure 4(a) shows the neighborhood 𝒩_90_ of residue 90, and Fig. 4(b) shows the weights (attention scores) of the edges in Fig. 4(b) using color gradient. Deeper hues indicate higher levels of attention. This residue is an interaction site and EGRET rightly predicted this. The other interaction sites, predicted by EGRET, in this neighborhood are shown in green. Interestingly, the edges with relatively higher attention scores (darker color hues) are mostly associated with source nodes that are predicted as interaction sites (residues at positions 50, 51, 52, 53, 55, 57, 59). Among these residues, 50, 52, 53, and 59 are true interaction sites (these four residues are associated to four of the top five attention scores). Moreover, the attention scores of the edges with non-interacting source nodes (e.g., 86, 87, 88, 89, 91, 92, 93) which are closer to 90 in primary sequence are lower than the attention scores of those associated with the long-range interactions (e.g., 50, 51, 52, 53, 59).

These results suggest that the patterns of attention scores, produced by EGRET, are not random, but are aligned with biological insights and carry meaningful information. This meaningful pattern of attention score provides insight into our model’s behaviour and decisions on interacting and non-interacting sites. While these results are promising, especially considering the black-box nature of other deep learning based methods, they should be interpreted with care. The interaction sites suggested by attention scores *alone* may contain false positives (e.g., residue 51) and false negatives. Higher attention scores do not necessarily guarantee an interaction site, nor is it certain that all the interaction sites within the neighborhood of another interaction site will have relatively higher attention scores. Indeed, the predictions of EGRET does not solely depend on the attention scores as it rightly predicted residue 58 and 89 to be non-interaction sites despite their associated edges having high attention scores. Followup studies are required to further investigate the interpretability of such graph based models as well as to design an attention mechanism so that the attention scores are more closely related to true interaction site predictions.

### 3.12 Running time

We investigated the time required to generate the features used by EGRET and DeepPPISP (one of most accurate structure-based methods) and the time required for prediction. All analyses were run on the same machine with Intel core i7-7700 CPU (4 cores), 16GB RAM, NVIDIA GeForce GTX 1070 GPU (8GB memory).

EGRET is much more faster than DeepPPISP. We report feature generation time on the smallest (39 amino acids) and the largest (500 amino acids) protein sequences in the test set (see Table 7). These results suggest that generating ProtBERT-based features is remarkably faster than generating the DeepPPISP features. For example, generating ProtBERT-based features took around only one minute for a 500 amino acid long protein, whereas it took around 2.5 hours for the DeepPPISP features since PSSM generation is more time-consuming. Moreover, PPI prediction time of EGRET, given the generated features, is also faster than DeepPPISP.

**Table 7:** Running time comparison between EGRET and DeepPPISP. We show the times (in seconds) required for generating the features and performing the prediction on the shortest and longest protein sequences in the test set.

| Protein length | Method | Only inference time | Feature extraction time | Total time |
| --- | --- | --- | --- | --- |
| 500 | EGRET | 4.54±0.05 | 63.28±0.1 | 67.82±0.15 |
|  | DeepPPISP | 15.02±0.2 | 8823.07±15 | 8838.09±15.2 |
| 39 | EGRET | 4.57±0.05 | 11.84±0.02 | 16.41±0.07 |
|  | DeepPPISP | 12.21±0.15 | 3706.74±6 | 3718.95±6.15 |

## 4 Conclusions

We have presented EGRET, a novel, highly accurate, and fast method for PPISP for isolated proteins. We have augmented GAT with edge aggregation and demonstrated its efficacy in improving the performance of PPISP. We also, for the first time, utilized transfer-learning with ProtBERT generated feature sets in PPISP. Our experimental results suggest that GAT (with or without edge aggregation) is better than other competing methods. We systematically analyzed the effects of our proposed edge aggregation and transfer learning with pretrained transformer-like models, and revealed that both of them have positive impact on PPISP. Furthermore, we investigated the performance of different methods under various model conditions with varying levels of long-range interactions and protein lengths, and demonstrated the superiority of our proposed methods. The demonstrated performance improvement across all seven evaluation metrics is quite promising. Thus, we believe EGRET advances the state-of-the-art in this domain, and will be considered as a useful tool for PPISP.

The use of edge aggregation to effectively capture the structural information is timely considering the growing evidence for the efficacy of structural information in residue-level prediction as well as the increasing availability of structure-known proteins. This study can be extended in several directions. We have adapted and systematically analyzed the ProtBERT model in Prot-Trans [43]. Follow-up studies will need to investigate other pretrained language models (e.g., Transformer-XL and XLNet) available in ProtTrans. Although we have focused on the PPI site prediction, EGRET has the potential to be investigated for the prediction of other types of binding sites, including DNA binding sites [75] and ligand (non-protein) binding sites [76]. Another potential research direction is to investigate the efficacy of EGRET and GAT-PPI on pair-wise PPI site prediction [30, 31] which we left as a future work.

## 5 Data Availability

The datasets underlying this article were derived from sources in the public domain. The benchmark DeepPPISP datasets were collected from https://github.com/CSUBioGroup/DeepPPISP. The MaSIF dataset was obtained from https://github.com/LPDI-EPFL/masif. The Dockground dataset is available at http://dockground.compbio.ku.edu/.

## References

[1] Javier De Las Rivas and Celia Fontanillo. Protein–protein interactions essentials: key concepts to building and analyzing interactome networks. PLoS Comput Biol, 6(6):e1000807, 2010.

[2] Naoki Orii and Madhavi K Ganapathiraju. Wiki-pi: a web-server of annotated human protein-protein interactions to aid in discovery of protein function. PloS one, 7(11):e49029, 2012.

[3] Khaled S Ahmed, Nahed H Saloma, and Yasser M Kadah. Improving the prediction of yeast protein function using weighted protein-protein interactions. Theoretical Biology and Medical Modelling, 8(1):11, 2011.

[4] Xingyi Li, Wenkai Li, Min Zeng, Ruiqing Zheng, and Min Li. Network-based methods for predicting essential genes or proteins: a survey. Briefings in bioinformatics, 21(2):566–583, 2020.

[5] Uros Kuzmanov and Andrew Emili. Protein-protein interaction networks: probing disease mechanisms using model systems. Genome medicine, 5(4):1–12, 2013.

[6] Rod K Nibbe, Salim A Chowdhury, Mehmet Koyutürk, Rob Ewing, and Mark R Chance. Protein–protein interaction networks and subnetworks in the biology of disease. Wiley Interdisciplinary Reviews: Systems Biology and Medicine, 3(3):357–367, 2011.

[7] Ioanna Petta, Sam Lievens, Claude Libert, Jan Tavernier, and Karolien De Bosscher. Modulation of protein–protein interactions for the development of novel therapeutics. Molecular Therapy, 24(4):707–718, 2016.

[8] Olivier Sperandio. Toward the design of drugs on protein-protein interactions. Current Pharmaceutical Design, 18(30):4585, 2012.

[9] Shoshana J Wodak, James Vlasblom, Andrei L Turinsky, and Shuye Pu. Protein–protein interaction networks: the puzzling riches. Current Opinion in Structural Biology, 23(6):941–953, 2013.

[10] Leandra M Brettner and Joanna Masel. Protein stickiness, rather than number of functional protein-protein interactions, predicts expression noise and plasticity in yeast. BMC Systems Biology, 6(1):128, 2012.

[11] AA Terentiev, NT Moldogazieva, and KV Shaitan. Dynamic proteomics in modeling of the living cell. protein-protein interactions. Biochemistry (Moscow), 74(13):1586–1607, 2009.

[12] Tobias Hamp and Burkhard Rost. More challenges for machine-learning protein interactions. Bioinformatics, 31(10):1521–1525, 2015.

[13] Iakes Ezkurdia, Lisa Bartoli, Piero Fariselli, Rita Casadio, Alfonso Valencia, and Michael L Tress. Progress and challenges in predicting protein–protein interaction sites. Briefings in Bioinformatics, 10(3):233–246, 2009.

[14] Loic Giot, Joel S Bader, C Brouwer, Amitabha Chaudhuri, Bing Kuang, Y Li, YL Hao, CE Ooi, Brian Godwin, E Vitols, et al. A protein interaction map of drosophila melanogaster. Science, 302(5651):1727–1736, 2003.

[15] Min Zeng, Fuhao Zhang, Fang-Xiang Wu, Yaohang Li, Jianxin Wang, and Min Li. Protein–protein interaction site prediction through combining local and global features with deep neural networks. Bioinformatics, 36(4):1114–1120, 2020.

[16] Thomas C Northey, Anja Barešić, and Andrew CR Martin. Intpred: a structure-based predictor of protein–protein interaction sites. Bioinformatics, 34(2):223–229, 2018.

[17] Tristan T Aumentado-Armstrong, Bogdan Istrate, and Robert A Murgita. Algorithmic approaches to protein-protein interaction site prediction. Algorithms for Molecular Biology, 10(1):7, 2015.

[18] Qingzhen Hou, Paul FG De Geest, Wim F Vranken, Jaap Heringa, and K Anton Feenstra. Seeing the trees through the forest: sequence-based homo-and heteromeric protein-protein interaction sites prediction using random forest. Bioinformatics, 33(10):1479–1487, 2017.

[19] Juan Fernandez-Recio, Maxim Totrov, and Ruben Abagyan. Identification of protein–protein interaction sites from docking energy landscapes. Journal of molecular biology, 335(3):843–865, 2004.

[20] Bowen Dai and Chris Bailey-Kellogg. Protein interaction interface region prediction by geometric deep learning. Bioinformatics, 2021.

[21] Aleksey Porollo and Jaroslaw Meller. Prediction-based fingerprints of protein–protein interactions. Proteins: Structure, Function, and Bioinformatics, 66(3):630–645, 2007.

[22] Huiling Chen and Huan-Xiang Zhou. Prediction of interface residues in protein–protein complexes by a consensus neural network method: test against nmr data. Proteins: Structure, Function, and Bioinformatics, 61(1):21–35, 2005.

[23] David La and Daisuke Kihara. A novel method for protein–protein interaction site prediction using phylogenetic substitution models. Proteins: Structure, Function, and Bioinformatics, 80(1):126–141, 2012.

[24] Pablo Gainza, Freyr Sverrisson, Frederico Monti, Emanuele Rodola, D Boscaini, MM Bronstein, and BE Correia. Deciphering interaction fingerprints from protein molecular surfaces using geometric deep learning. Nature Methods, 17(2):184–192, 2020.

[25] Yiwei Li, G Brian Golding, and Lucian Ilie. DELPHI: accurate deep ensemble model for protein interaction sites prediction. Bioinformatics, 08 2020. btaa750.

[26] Jian Zhang and Lukasz Kurgan. Scriber: accurate and partner type-specific prediction of protein-binding residues from proteins sequences. Bioinformatics, 35(14):i343–i353, 2019.

[27] Xiaoying Wang, Bin Yu, Anjun Ma, Cheng Chen, Bingqiang Liu, and Qin Ma. Protein–protein interaction sites prediction by ensemble random forests with synthetic minority oversampling technique. Bioinformatics, 35(14):2395–2402, 2019.

[28] G Singh, K Dhole, PP Pai, and S Mondal. Springs: Prediction of protein-protein interaction sites using artificial neural networks. J Proteomics Computational Biol, 1(1):7, 2014.

[29] Yoichi Murakami and Kenji Mizuguchi. Applying the naïve bayes classifier with kernel density estimation to the prediction of protein–protein interaction sites. Bioinformatics, 26(15):1841–1848, 2010.

[30] Alex Fout, Jonathon Byrd, Basir Shariat, and Asa Ben-Hur. Protein interface prediction using graph convolutional networks. In Advances in neural information processing systems, pages 6530–6539, 2017.

[31] Raphael Townshend, Rishi Bedi, Patricia Suriana, and Ron Dror. End-to-end learning on 3d protein structure for interface prediction. In Advances in Neural Information Processing Systems, pages 15642–15651, 2019.

[32] Ruben Sanchez-Garcia, Carlos Oscar Sánchez Sorzano, José María Carazo, and Joan Segura. Bipspi: a method for the prediction of partner-specific protein–protein interfaces. Bioinformatics, 35(3):470–477, 2019.

[33] Susan Jones and Janet M Thornton. Analysis of protein-protein interaction sites using surface patches. Journal of molecular biology, 272(1):121–132, 1997.

[34] Shandar Ahmad and Kenji Mizuguchi. Partner-aware prediction of interacting residues in protein-protein complexes from sequence data. PLoS One, 6(12):e29104, 2011.

[35] Zhi-Sen Wei, Ke Han, Jing-Yu Yang, Hong-Bin Shen, and Dong-Jun Yu. Protein–protein interaction sites prediction by ensembling svm and sample-weighted random forests. Neurocomputing, 193:201–212, 2016.

[36] Yanay Ofran and Burkhard Rost. Isis: interaction sites identified from sequence. Bioinformatics, 23(2):e13–e16, 2007.

[37] Changhui Yan, Drena Dobbs, and Vasant Honavar. A two-stage classifier for identification of protein–protein interface residues. Bioinformatics, 20(Suppl 1):i371–i378, 2004.

[38] Xiaoying Wang, Bin Yu, Anjun Ma, Cheng Chen, Bingqiang Liu, and Qin Ma. Protein–protein interaction sites prediction by ensemble random forests with synthetic minority oversampling technique. Bioinformatics, 35(14):2395–2402, 2019.

[39] Josip Mihel, Mile Šikić, Sanja Tomić, Branko Jeren, and Kristian Vlahoviček. Psaia–protein structure and interaction analyzer. BMC Structural Biology, 8(1):21, 2008.

[40] Mostofa Rafid Uddin, Sazan Mahbub, M Saifur Rahman, and Md Shamsuzzoha Bayzid. SAINT: self-attention augmented inception-inside-inception network improves protein secondary structure prediction. Bioinformatics, 36(17):4599–4608, 2020.

[41] Jack Hanson, Kuldip Paliwal, Thomas Litfin, Yuedong Yang, and Yaoqi Zhou. Improving prediction of protein secondary structure, backbone angles, solvent accessibility and contact numbers by using predicted contact maps and an ensemble of recurrent and residual convolutional neural networks. Bioinformatics, 35(14):2403–2410, 2019.

[42] Stephen F Altschul, Thomas L Madden, Alejandro A Schäffer, Jinghui Zhang, Zheng Zhang, Webb Miller, and David J Lipman. Gapped blast and psi-blast: a new generation of protein database search programs. Nucleic Acids Research, 25(17):3389–3402, 1997.

[43] Ahmed Elnaggar, Michael Heinzinger, Christian Dallago, Ghalia Rihawi, Yu Wang, Llion Jones, Tom Gibbs, Tamas Feher, Christoph Angerer, Debsindhu Bhowmik, et al. Prottrans: Towards cracking the language of life’s code through self-supervised deep learning and high performance computing. arXiv preprint 2007.06225, 2020.

[44] Zihang Dai, Zhilin Yang, Yiming Yang, Jaime G Carbonell, Quoc Le, and Ruslan Salakhutdinov. Transformer-xl: Attentive language models beyond a fixed-length context. In Proceedings of the 57th Annual Meeting of the Association for Computational Linguistics, pages 2978–2988, 2019.

[45] Zhilin Yang, Zihang Dai, Yiming Yang, Jaime Carbonell, Russ R Salakhutdinov, and Quoc V Le. Xlnet: Generalized autoregressive pretraining for language understanding. In Advances in neural information processing systems, pages 5753–5763, 2019.

[46] Jacob Devlin, Ming-Wei Chang, Kenton Lee, and Kristina Toutanova. BERT: Pre-training of deep bidirectional transformers for language understanding. In Proceedings of the 2019 Conference of the North American Chapter of the Association for Computational Linguistics: Human Language Technologies, Volume 1 (Long and Short Papers), pages 4171–4186, Minneapolis, Minnesota, June 2019. Association for Computational Linguistics.

[47] Zhenzhong Lan, Mingda Chen, Sebastian Goodman, Kevin Gimpel, Piyush Sharma, and Radu Soricut. Albert: A lite bert for self-supervised learning of language representations. In International Conference on Learning Representations, 2019.

[48] Jesse Vig, Ali Madani, Lav R Varshney, Caiming Xiong, Richard Socher, and Nazneen Fatema Rajani. Bertology meets biology: Interpreting attention in protein language models. arXiv preprint 2006.15222, 2020.

[49] Buzhong Zhang, Jinyan Li, Lijun Quan, Yu Chen, and Qiang Lü. Sequence-based prediction of protein-protein interaction sites by simplified long short-term memory network. Neurocomputing, 357:86–100, 2019.

[50] Helen M Berman, John Westbrook, Zukang Feng, Gary Gilliland, Talapady N Bhat, Helge Weissig, Ilya N Shindyalov, and Philip E Bourne. The protein data bank. Nucleic Acids Research, 28(1):235–242, 2000.

[51] Thomas N. Kipf and Max Welling. Semi-supervised classification with graph convolutional networks. In International Conference on Learning Representations (ICLR), 2017.

[52] Petar Veličković, Guillem Cucurull, Arantxa Casanova, Adriana Romero, Pietro Lío, and Yoshua Bengio. Graph attention networks. In International Conference on Learning Representations, 2018.

[53] Yue Wang, Yongbin Sun, Ziwei Liu, Sanjay E Sarma, Michael M Bronstein, and Justin M Solomon. Dynamic graph cnn for learning on point clouds. Acm Transactions On Graphics (tog), 38(5):1–12, 2019.

[54] Yi Liu, Hao Yuan, Lei Cai, and Shuiwang Ji. Deep learning of high-order interactions for protein interface prediction. In Proceedings of the 26th ACM SIGKDD International Conference on Knowledge Discovery & Data Mining, pages 679–687, 2020.

[55] Dzmitry Bahdanau, Kyunghyun Cho, and Yoshua Bengio. Neural machine translation by jointly learning to align and translate. In 3rd International Conference on Learning Representations, ICLR 2015, 2015.

[56] Ashish Vaswani, Noam Shazeer, Niki Parmar, Jakob Uszkoreit, Llion Jones, Aidan N Gomez, Lukasz Kaiser, and Illia Polosukhin. Attention is all you need. In Advances in neural information processing systems, pages 5998–6008, 2017.

[57] Peng Wang, Qi Wu, Jiewei Cao, Chunhua Shen, Lianli Gao, and Anton van den Hengel. Neighbourhood watch: Referring expression comprehension via language-guided graph attention networks. In Proceedings of the IEEE/CVF Conference on Computer Vision and Pattern Recognition, pages 1960–1968, 2019.

[58] David Eppstein, Michael S Paterson, and F Frances Yao. On nearest-neighbor graphs. Discrete & Computational Geometry, 17(3):263–282, 1997.

[59] IUPAC-IUB Tentative Rules. A one letter notation for amino acid sequence. Biochem. J, 113:1–4, 1969.

[60] Marinka Zitnik and Jure Leskovec. Predicting multicellular function through multi-layer tissue networks. Bioinformatics, 33(14):i190–i198, 2017.

[61] Jun Han and Claudio Moraga. The influence of the sigmoid function parameters on the speed of backpropagation learning. In International Workshop on Artificial Neural Networks, pages 195–201. Springer, 1995.

[62] Minjie Wang, Lingfan Yu, D. Zheng, Quan Gan, Yu Gai, Zihao Ye, Mufei Li, Jinjing Zhou, Qi Huang, Chao Ma, et al. Deep graph library: Towards efficient and scalable deep learning on graphs. arXiv preprint 1909.01315, 2019.

[63] Howook Hwang, Brian Pierce, Julian Mintseris, Jöel Janin, and Zhiping Weng. Protein–protein docking benchmark version 3.0. Proteins: Structure, Function, and Bioinformatics, 73(3):705–709, 2008.

[64] Utkan Ogmen, Ozlem Keskin, A Selim Aytuna, Ruth Nussinov, and Attila Gursoy. Prism: protein interactions by structural matching. Nucleic acids research, 33(Suppl 2):W331–W336, 2005.

[65] Alper Baspinar, Engin Cukuroglu, Ruth Nussinov, Ozlem Keskin, and Attila Gursoy. Prism: a web server and repository for prediction of protein–protein interactions and modeling their 3d complexes. Nucleic acids research, 42(W1):W285–W289, 2014.

[66] Brian G Pierce, Kevin Wiehe, Howook Hwang, Bong-Hyun Kim, Thom Vreven, and Zhiping Weng. Zdock server: interactive docking prediction of protein–protein complexes and symmetric multimers. Bioinformatics, 30(12):1771–1773, 2014.

[67] Renxiao Wang, Xueliang Fang, Yipin Lu, Chao-Yie Yang, and Shaomeng Wang. The pdbbind database: methodologies and updates. Journal of medicinal chemistry, 48(12):4111–4119, 2005.

[68] James Dunbar, Konrad Krawczyk, Jinwoo Leem, Terry Baker, Angelika Fuchs, Guy Georges, Jiye Shi, and Charlotte M Deane. Sabdab: the structural antibody database. Nucleic acids research, 42(D1):D1140–D1146, 2014.

[69] Ying Huang, Beifang Niu, Ying Gao, Limin Fu, and Weizhong Li. Cd-hit suite: a web server for clustering and comparing biological sequences. Bioinformatics, 26(5):680–682, 2010.

[70] Petras J Kundrotas, Ivan Anishchenko, Taras Dauzhenka, Ian Kotthoff, Daniil Mnevets, Matthew M Copeland, and Ilya A Vakser. Dockground: a comprehensive data resource for modeling of protein complexes. Protein Science, 27(1):172–181, 2018.

[71] Sjoerd J de Vries and Alexandre MJJ Bonvin. How proteins get in touch: interface prediction in the study of biomolecular complexes. Current Protein and Peptide Science, 9(4):394–406, 2008.

[72] Min Zeng, Beiji Zou, Faran Wei, Xiyao Liu, and Lei Wang. Effective prediction of three common diseases by combining smote with tomek links technique for imbalanced medical data. In 2016 IEEE International Conference of Online Analysis and Computing Science (ICOACS), pages 225–228. IEEE, 2016.

[73] Frank Wilcoxon, SK Katti, and Roberta A Wilcox. Critical values and probability levels for the wilcoxon rank sum test and the wilcoxon signed rank test. Selected tables in mathematical statistics, 1:171–259, 1970.

[74] Rhys Heffernan, Yuedong Yang, Kuldip Paliwal, and Yaoqi Zhou. Capturing non-local interactions by long short-term memory bidirectional recurrent neural networks for improving prediction of protein secondary structure, backbone angles, contact numbers and solvent accessibility. Bioinformatics, 33(18):2842–2849, 2017.

[75] Shandar Ahmad and Akinori Sarai. Pssm-based prediction of dna binding sites in proteins. BMC bioinformatics, 6(1):33, 2005.

[76] Alasdair TR Laurie and Richard M Jackson. Q-sitefinder: an energy-based method for the prediction of protein–ligand binding sites. Bioinformatics, 21(9):1908–1916, 2005.

