## supplementary materials for "EGRET: Edge Aggregated Graph Attention Networks and Transfer Learning Improve Protein-Protein Interaction Site Prediction"

#### 1 Overview

These supplementary materials present additional details about the methods used (Secs. 2-3) and additional results (Secs. 4-6).

#### 2 Methods

Table S1 shows different types of features used by various methods considered in this study. SPPIDER [1] showed the impact of a combination of putative relative solvent accessibility (RSA) prediction-based fingerprints with other structural and sequence information using Support Vector Machines (SVM) and neural networks. IntPred [2] exploited the random forest algorithm and used various structural features as well as sequence-based features. DeepPPISP [3] is a deep learning based method that used both local and global features generated from eight-state secondary structures (derived from PDB [10] structure files using DSSP [11, 12][1] program), and various sequence-based features (raw protein sequence and position-specific score matrices (PSSM)). DELPHI [4] – one the most accurate methods – used an ensemble of two deep learning models (one convolutional based and one bidirectional Gated Recurrent Unit [13] based neural network) as the predictor, and was trained in an ensemble learning fashion using a dataset of 9,982 protein sequences, where each residue is represented by twelve types of features (see Table S1 for more details). SCIBBER [5] used a training set comprising 843 proteins containing several primary sequence-based features and their derivatives (e.g., Relative Amino Acid Propensity (RAAP) for binding, Putative Relative Solvent Accessibility (RSA), Hydrophobicity,

[1]https://swift.cmbi.umcn.nl/gv/dssp/

Table S1: Feature sets used by various state-of-the-art methods.

| **Method** | **Feature-type** | **Features** |
|---|---|---|
| SPPIDER [1] | Sequence- and structure-based | Putative relative solvent accessibility (RSA), Surface patch of protein |
| IntPred [2] | Sequence- and structure-based | Disulphide bonds, hydrogen bonds and secondary structures ($\alpha$-helix, $\beta$-sheet, mixed secondary structure, and coil), planarity, propensity score, hydrophobicity, homology conservation score and FEP conservation score |
| DeepPPISP [3] | Sequence- and Structure-based | Eight-state secondary structures, raw protein sequence and position-specific score matrices (PSSM) |
| DELPHI [4] | Sequence-based | High-scoring segment pair (HSP), a variation of 3-mer amino acid embedding (ProtVec1D), position information, position-specific scoring matrix (PSSM), evolutionary conservation (ECO), putative relative solvent accessibility (RSA), relative amino acid propensity (RAA), putative protein-binding disorder, hydropathy index, physicochemical characteristics, physical properties, and PKx |
| SCIBBER [5] | Sequence-based | Relative Amino Acid Propensity (RAAP) for binding, Putative Relative Solvent Accessibility (RSA), evolutionary conservation (ECO), Hydrophobicity, Polarity, Charge, protein-binding disordered regions, putative secondary structure, physicochemical properties, relative position, relative position |
| PSIVER [6] | Sequence-based | Predicted accessibility and PSSM |
| SPRINGS [7] | Sequence-based | Evolutionary information, averaged cumulative hydropathy and predicted RSA |
| ISIS [8] | Sequence-based | predicted structural features, evolutionary information |
| RF_PPI [9] | Sequence-based | Protein size, predicted backbone flexibility, sequence specificity |

etc.), and predicted interacting residues using a two-layer design, where the first and second layers consist of five and one machine learning models, respectively. PSIVER [6] trained a Naïve Bayes classifier on sequence-based features (predicted accessibility and PSSM). SPRINGS [7] proposed a shallow artificial neural network, which uses various sequence based features (e.g., evolutionary information, averaged cumulative hydropathy and predicted RSA). ISIS [8] used shallow neural networks and combined predicted structural features with evolutionary information as feature representations. RF_PPI [9] presented a study on the evaluation of the importance of various sequence based features using Random Forest (RF) classifier.

### 3 Evaluation metrics

$$Accuracy = \frac{TP + TN}{TP + TN + FP + FN} \tag{1}$$

$$Precision = \frac{TP}{TP + FP} \tag{2}$$

$$Recall = \frac{TP}{TP + FN} \tag{3}$$

$$F1 - measure = \frac{2 * Precision * Recall}{Precision + Recall} \tag{4}$$

$$MCC = \frac{TP * TN - FP * FN}{\sqrt{(TP + FP)(TP + FN)(TN + FP)(TN + FN)}} \tag{5}$$

In Eqns. 1-5, true positives (TP) and true negatives (TN) represent the number of correctly predicted interaction and non-interaction sites respectively. Similarly, the number of incorrectly predicted interaction and non-interaction sites are represented by false positives (FP) and false negatives (FN), respectively.

### 4 Statistical significance test

Table S2: The statistical significance of various performance metrics between different pairs of methods. We show the $p$-values using the Wilcoxon signed rank test.

| Method-pair | F-measure | AUROC | AUPRC | MCC |
|---|---|---|---|---|
| DELPHI, EGRET | 0.0042 | 0.0674 | 0.0457 | 0.0079 |
| DELPHI, GAT-PPI | 0.0153 | 0.0535 | 0.0369 | 0.0372 |
| GAT-PPI, EGRET | 0.0719 | 0.1861 | 0.2139 | 0.1057 |

### 5 Impact of long-range interactions

Table S3: AUPRC of EGRET and DELPHI under various levels of non-local interactions. 70 proteins in the testset were divided into 7 disjoint bins each having 10 proteins. The binning was based on the number of non-local contacts per residue in the proteins.

| Non-local contact per residue | AUPRC of EGRET | AUPRC of DELPHI | AUPRC Difference | $p$-value |
|---|---|---|---|---|
| 0.47-1.49 | 0.5666 | 0.5787 | -0.0121 | 0.8389 |
| 1.54-1.72 | 0.5243 | 0.4733 | 0.0510 | 0.4062 |
| 1.73-1.93 | 0.4800 | 0.4321 | 0.0479 | 0.2929 |
| 1.93-2.15 | 0.4461 | 0.4044 | 0.0417 | 0.4044 |
| 2.16-2.28 | 0.4030 | 0.3765 | 0.0265 | 0.3515 |
| 2.29-2.51 | 0.3953 | 0.3660 | 0.0293 | 0.1446 |
| 2.59-3.21 | 0.4046 | 0.3600 | 0.0446 | 0.0457 |

Table S4: AUPRC of GAT-PPI and DELPHI under various levels of non-local interactions. 70 proteins in the testset were divided into 7 disjoint bins each having 10 proteins. The binning was based on the number of non-local contacts per residue in the proteins.

| Non-local contact per residue | AUPRC of GAT-PPI | AUPRC of DELPHI | AUPRC Difference | $p$-value |
|---|---|---|---|---|
| 0.47-1.49 | 0.5318 | 0.5787 | -0.0469 | 0.8838 |
| 1.54-1.72 | 0.4972 | 0.4733 | 0.0239 | 0.5218 |
| 1.73-1.93 | 0.4537 | 0.4321 | 0.0215 | 0.4959 |
| 1.93-2.15 | 0.4288 | 0.4044 | 0.0244 | 0.3940 |
| 2.16-2.28 | 0.4089 | 0.3765 | 0.0324 | 0.2764 |
| 2.29-2.51 | 0.3966 | 0.3660 | 0.0306 | 0.1602 |
| 2.59-3.21 | 0.3937 | 0.3600 | 0.0338 | 0.0369 |

### 6 Impact of protein length

Table S5: AUPRC of EGRET and DELPHI under various levels of protein lengths. 70 proteins in the testset were divided into 7 disjoint bins each having 10 proteins. The binning was based on the lengths of the proteins.

| Protein Length | AUPRC of EGRET | AUPRC of DELPHI | AUPRC Difference | $p$-value |
|---|---|---|---|---|
| 51-77 | 0.5937 | 0.5474 | 0.0463 | 0.7539 |
| 82-102 | 0.5293 | 0.4795 | 0.0499 | 0.5796 |
| 104-131 | 0.4813 | 0.4547 | 0.0266 | 0.6856 |
| 144-165 | 0.4697 | 0.4231 | 0.0466 | 0.2726 |
| 169-198 | 0.4453 | 0.3991 | 0.0461 | 0.1204 |
| 205-273 | 0.4331 | 0.3843 | 0.0488 | 0.0374 |
| 304-376 | 0.4046 | 0.3600 | 0.0446 | 0.0457 |

Table S6: AUPRC of GAT-PPI and DELPHI under various levels of protein lengths. 70 proteins in the testset were divided into 7 disjoint bins each having 10 proteins. The binning was based on the lengths of the proteins.

| Protein Length | AUPRC of GAT-PPI | AUPRC of DELPHI | AUPRC Difference | $p$-value |
|---|---|---|---|---|
| 51-77 | 0.5879 | 0.5474 | 0.0405 | 0.5391 |
| 82-102 | 0.5132 | 0.4795 | 0.0337 | 0.3506 |
| 104-131 | 0.4756 | 0.4547 | 0.0209 | 0.5287 |
| 144-165 | 0.4572 | 0.4231 | 0.0341 | 0.1666 |
| 169-198 | 0.4390 | 0.3991 | 0.0399 | 0.1056 |
| 205-273 | 0.4233 | 0.3843 | 0.0390 | 0.0503 |
| 304-376 | 0.3937 | 0.3600 | 0.0338 | 0.0369 |

### References


[1] Aleksey Porollo and Jarosław Meller. Prediction-based fingerprints of protein–protein interactions. *Proteins: Structure, Function, and Bioinformatics*, 66(3):630–645, 2007.

[2] Thomas C Northey, Anja Barešić, and Andrew CR Martin. Intpred: a structure-based predictor of protein–protein interaction sites. *Bioinformatics*, 34(2):223–229, 2018.

[3] Min Zeng, Fuhao Zhang, Fang-Xiang Wu, Yaohang Li, Jianxin Wang, and Min Li. Protein–protein interaction site prediction through combining local and global features with deep neural networks. *Bioinformatics*, 36(4):1114–1120, 2020.

[4] Yiwei Li, G Brian Golding, and Lucian Ilie. DELPHI: accurate deep ensemble model for protein interaction sites prediction. *Bioinformatics*, 08 2020. btaa750.

[5] Jian Zhang and Lukasz Kurgan. Scriber: accurate and partner type-specific prediction of protein-binding residues from proteins sequences. *Bioinformatics*, 35(14):i343–i353, 2019.

[6] Yoichi Murakami and Kenji Mizuguchi. Applying the naïve bayes classifier with kernel density estimation to the prediction of protein–protein interaction sites. *Bioinformatics*, 26(15):1841–1848, 2010.

[7] G Singh, K Dhole, PP Pai, and S Mondal. Springs: Prediction of protein-protein interaction sites using artificial neural networks. *J Proteomics Computational Biol*, 1(1):7, 2014.

[8] Yanay Ofran and Burkhard Rost. Isis: interaction sites identified from sequence. *Bioinformatics*, 23(2):e13–e16, 2007.

[9] Qingzhen Hou, Paul FG De Geest, Wim F Vranken, Jaap Heringa, and K Anton Feenstra. Seeing the trees through the forest: sequence-based homo-and heteromeric protein-protein interaction sites prediction using random forest. *Bioinformatics*, 33(10):1479–1487, 2017.

[10] Helen M Berman, John Westbrook, Zukang Feng, Gary Gilliland, Talapady N Bhat, Helge Weissig, Ilya N Shindyalov, and Philip E Bourne. The protein data bank. *Nucleic Acids Research*, 28(1):235–242, 2000.

[11] Wouter G Touw, Coos Baakman, Jon Black, Tim AH Te Beek, Elmar Krieger, Robbie P Joosten, and Gert Vriend. A series of pdb-related databanks for everyday needs. *Nucleic Acids Research*, 43(D1):D364–D368, 2015.

[12] Wolfgang Kabsch and Christian Sander. Dictionary of protein secondary structure: pattern recognition of hydrogen-bonded and geometrical features. *Biopolymers: Original Research on Biomolecules*, 22(12):2577–2637, 1983.

[13] Kyunghyun Cho, Bart van Merrienboer, Çaglar Gülçehre, Dzmitry Bahdanau, Fethi Bougares, Holger Schwenk, and Yoshua Bengio. Learning phrase representations using rnn encoder-decoder for statistical machine translation. In *EMNLP*, 2014.
